# Gastric resection and restriction yield distinct neuroendocrine and metabolic outcomes in obese mice

**DOI:** 10.64898/2026.10.27.762188

**Authors:** Mohammad Jarrah, Sanaz Saleh, Dana Tasabehji, Aviva Fraer, Yi Chu, Zobayda Farzana Haque, Andrew C Shin, Mohamad Mokadem

**Author notes:** Corresponding author & Lead contact.

## Abstract

Whether the metabolic benefits of sleeve gastrectomy arise from gastric restriction alone or also depend on tissue resection remains unclear. We compared sleeve gastrectomy (SG) with a surgical clip-based sleeve gastroplasty (GP) model that restricts the stomach without removing tissue. Diet-induced obese male C57BL/6J mice were randomized to SG, GP, or sham surgery and evaluated over six weeks for energy balance, glucose regulation, hepatic lipid accumulation, circulating hormones and branched-chain amino acids (BCAAs), and hypothalamic neuropeptide expression. SG and GP produced comparable weight loss of 12% and 11%, respectively, without significant differences in energy expenditure. However, the procedures differed in the timing of reduced food intake, and only SG produced smaller, more frequent meals. Both procedures improved glucose handling, with greater improvement after SG. SG also selectively reduced hepatic triglyceride content and circulating BCAAs. Hypothalamic POMC expression decreased after both procedures, whereas NPY and AgRP expression decreased only after SG. SG further produced a distinct hormonal profile, including higher fasting GLP-1 and PYY, lower ghrelin, and greater glucose-stimulated increases in several gut and pancreatic hormones. Thus, comparable short-term weight loss after gastric resection and non-resection restriction was accompanied by distinct metabolic and neuroendocrine responses. These findings support a contribution of gastric resection beyond restriction alone and identify candidate gut–brain and metabolic pathways for developing less invasive interventions that reproduce the broader benefits of sleeve gastrectomy.

## Introduction

Obesity remains a major global health challenge, affecting over 650 million individuals worldwide and contributing to millions of deaths annually due to associated complications such as heart disease and stroke [1, 2]. It is an alarming issue in the United States of America, where predictions suggest that 50% of the population may have obesity by 2030 [3]. Bariatric surgeries have emerged as durable and effective treatments for obesity, offering improvements in obesity-related morbidity and reduced mortality compared to non-surgical interventions [4-7]. The two primary surgical interventions, Roux-en-Y gastric bypass (RYGB) and sleeve gastrectomy (SG), are commonly performed as bariatric surgeries [4]. SG, introduced in 1990, has become the most prevalent bariatric procedure in the United States over the past two decades [8]. On the other hand, bariatric endoscopic procedures are interventions that replicate bariatric surgeries in a less invasive manner to achieve a considerable amount of weight loss and improve the metabolic profile of patients with obesity [1]. Endoscopic sleeve gastroplasty (ESG) presents a less invasive alternative to sleeve gastrectomy, employing a restrictive mechanism to reduce stomach size without tissue removal. It involves using a suturing device to create opposing sutures inside the stomach, restricting a large portion of the stomach without resection, excluding contact of food with the gastric body, and inducing an early sensation of fullness [9]. Although ESG is a relatively new procedure, several studies have proved it to be a safe and effective intervention for weight loss [10-12]. We have replicated bariatric surgeries such as SG and RYGB in mice and have examined the metabolic improvements these surgeries provide [13, 14]. Other studies investigated the diverse (and weight-independent) advantages of SG on glucose metabolism [15] and liver steatosis [16, 17] in mice. However, it is still not known whether the metabolic effects of this procedure are due to the removal of that portion of the stomach or the actual exclusion of food contact with it. Therefore, we developed two surgical models of SG and clip-based sleeve gastroplasty (GP) in mice, where we restrict the gastric body in a sleeve shape using a clip and sutures, and then we either resect (in SG) or keep that part of the stomach intact (in GP). Although this GP model is not an endoscopic sleeve gastroplasty (ESG) and does not replicate the endoscopic suturing technique used clinically; it represents a surgical-based, restrictive gastroplasty model that preserves the gastric fundus and greater-curvature segment. Then we planned our study to characterize and comprehensively examine a battery of metabolic parameters between the two interventions compared to a sham procedure.

## Methods

### Animal care

Animal care and procedures were approved by the Institutional Animal Care and Use Committee (IACUC) of the University of Iowa. All animals were humanely euthanized at the designated experimental endpoints using CO₂ chambers, followed by exsanguination as a secondary physical method to ensure death, in accordance with our approved institutional animal care protocol and IACUC guidelines. All mice were housed in standard ventilated cages (maximum 4 mice per cage before surgery; single-housed only during designated food-intake measurements) with corn-cob bedding, nesting material, and a plastic shelter as environmental enrichment. Animals were maintained in a specific pathogen-free facility at 22–24°C with 40–60% humidity on a 12-hour light–dark cycle (lights on at 6:00). Animals had ad libitum access to high-fat diet and water except during defined fasting periods for metabolic testing. Health status and welfare were monitored at least once daily, and during the immediate postoperative period animals were checked a minimum of twice daily for hydration status, activity, incision integrity, and signs of pain or distress. CO₂ euthanasia followed by exsanguination was selected in accordance with American Veterinary Medical Association (AVMA) guidelines and was approved in the IACUC protocol as a humane, rapid method appropriate for adult mice.

### Study Design

Sixteen-week-old diet-induced obesity (DIO) C57BL/6J male mice were purchased from Jackson Laboratory (Bar Harbor, ME, USA). They were acclimated for two weeks on a high-fat diet (60% calories from fat; D12492, Research Diets) and water ad libitum. Animals were randomly allocated to SG, clip-based sleeve gastroplasty (GP), or sham surgery using a computer-generated randomization sequence (Excel RAND function), stratified by baseline body weight to ensure comparable mean weights across groups. The individual mouse was considered the experimental unit for all analyses, including body weight, food intake (averaged per animal over the measurement interval), energy expenditure, body composition, hormones, gene expression, BCAAs, and histologic quantification (where multiple fields per section were averaged to a single value per mouse).

Inclusion criteria were successful completion of the assigned surgery, recovery to normal activity and grooming within 48 hours, and maintenance on the study diet without signs of unrelated illness. Pre-specified exclusion criteria included intraoperative technical failure, perioperative complications requiring early humane euthanasia, or insufficient biospecimen volume or assay failure for specific measurements. For each endpoint, the exact number of animals analyzed (n) per group is reported in the figures and legends; any animal or data points excluded from a given analysis and the reasons are described in the Results or figure legends.

### Surgical Interventions

On the day of the intervention, all animals were given a weight-based dose of enrofloxacin (10 mg/Kg, subcutaneous; Baytril, Bayer) for perioperative infection prophylaxis and buprenorphine (0.05–0.1 mg/kg, subcutaneous; Buprenex, Reckitt Benckiser) for analgesia, administered 30 minutes before surgery and every 12 hours for 48 hours postoperatively, with additional doses as needed based on clinical assessment. General anesthesia was induced and maintained with isoflurane (1.5–3% in oxygen via nose cone)After removing hair with an electric razor, the abdomen skin was sterilized by 3 alternating scrubbings with betadine and 70% alcohol, then wiped with a betadine prep pad. A midline laparotomy was performed using forceps and scissors and extended from the xiphoid to the umbilicus. The abdominal wall was entered using forceps and sterile cotton swabs.

The SG procedure was replicated as previously described[13].A midline laparotomy was performed using forceps and scissors and extended from the xiphoid to the umbilicus. The abdominal wall was entered using forceps and sterile cotton swabs. For SG, the stomach was exteriorized, and the short gastric vessels were divided along the greater curvature, then a tubular gastric “sleeve” was created along the lesser curvature and the excluded greater-curvature stomach (including most of the fundus) was resected with scissors as previously described. GP followed the same initial steps of exposure and clip placement, but after the stomach was narrowed in a sleeve-like configuration using an external clip along the lesser curvature and supporting seromuscular sutures, the greater-curvature portion was left intact and not transected or removed. Thus, GP creates a restricted gastric channel while preserving the excluded greater-curvature segment and fundus in continuity, in contrast to SG where this segment is resected.. Closure was performed using 4−0 Vicryl with a running suture in the abdominal wall and interrupted sutures using 4−0 nylon. In the sham procedures, the abdomen was opened in the same manner, and a clip applicator was applied with gentle pressure to the outside of the stomach for several minutes along the line of resection of the sleeve. The bowels were manipulated without any rearrangement until the anesthesia time matched that of the actual surgery. Abdominal wall and skin closures were performed in the same manner. All mice recovered from the stress of surgery in their home cages with prophylactic injection of analgesics daily for the first 2 days and then on an as-needed basis. Mice were placed on a liquid diet and supplemental intraperitoneal injections of dextrose and fluid for hydration and nutrition during postoperative week 1. They were all allowed to recover on a solid diet during postoperative week 2. Body weight was measured weekly.

### Food intake and energy expenditure measurements

Food intake was measured during week 1, week 2 and week 3 post-intervention in single-housed mice over 5 consecutive days in each corresponding week. Daily food intake (kcal/day) was calculated as the difference between weighed food offered and remaining food (including spillage collected from the cage) divided by the number of days and averaged per mouse for each week. Feed efficiency was calculated as the change in body weight (g) divided by cumulative caloric intake (kcal) over the same time interval. Energy expenditure and food intake behavior, including meal size and frequency, were measured in Promethion metabolic cages (Sable Systems) by the University of Iowa Metabolic Phenotyping Core facility during week 4 post-intervention. Mice were acclimated to the metabolic cages for 24 hours before data collection; VO₂, VCO₂, locomotor activity, and food intake were then recorded continuously for at least 48 hours at 20–22°C on the same 12-hour light–dark cycle. Total energy expenditure (TEE) was derived from VO₂ and VCO₂ using the manufacturer’s software

### Glucose tolerance test (GTT)

GTT was performed following an overnight fast for 16 hours (postoperative week 4). Mice received 2 g/kg D-glucose intraperitoneal injection (cat no: 50–99-7, RPI, Mt. Prospect, IL). Blood glucose was measured from tail vein blood using a handheld glucometer (Contour, Bayer HealthCare LLC) immediately before and 15, 30, 45, 60, and 120 minutes after glucose administration.

### Insulin tolerance test (ITT)

ITT was performed following a 4-hour fast (postoperative week 5). Mice received 0.5 U/kg insulin (NDC 0002–8215-01, HI-210, Humulin R, Eli Lily) by intraperitoneal injection. Blood glucose was measured from tail vein blood using a handheld glucometer (Contour, Bayer HealthCare LLC) immediately before and 15, 30, 45, 60, and 120 minutes after insulin administration.

### Measurements of gut hormones

Baseline blood from SG, GP, and their sham counterparts was collected after an overnight fast retro-orbitally with Ethylenediaminetetraacetic acid (EDTA) coated Natelson blood collecting tube (02-668-10, Fisher Scientific). Blood was collected into a 1.5 ml Eppendorf tube containing protease inhibitor mix (Roche) and Dipeptidyl peptidase-4 (DPP4) inhibitor (DPP4, Millipore). The tubes were inverted several times to mix. A second blood collection was performed 10 minutes after oral gavage of 1 g/kg D-Glucose. Plasma was collected after centrifugation at 1,000x g for 10 minutes and stored at -80^0^C. The hormonal levels are presented as "baseline" (pre-gavage) and "stimulated" (post-gavage) measurements. A Milliplex mouse metabolic hormone expanded panel kit (MMHE-44K, Millipore) was used to determine plasma concentrations of the reported hormones. Steps were taken following the detailed instructions of the manufacturer. Bio-Plex 200 (Bio-Rad) was used for measurements with standard curves for each hormone. Total glucagon-like peptide-1 (GLP-1), insulin, C-peptide, glucagon, amylin, gastric inhibitory polypeptide (GIP), Peptide-YY (PYY), ghrelin, and leptin were measured in two serum samples: one after 12 hours of fasting and one after oral gavage of glucose (1g/Kg).

### Liver triglyceride (TG) measurement

Liver samples of around 200 mg were used for tissue levels of TG, using a triglyceride colorimetric assay kit (10010303, Cayman Chemical). Briefly, the tissue was homogenized and centrifuged. Insoluble fat on the top and all supernatants were further extracted for TG. Portions containing TG were mixed with an enzyme mix. The final product, quinoneimine, which reflects TG level, produces a brilliant purple color, which was measured at 540 nm. With TG standards, concentrations of TG were obtained. Final data were expressed as TG per mg of initial liver sample.

### Hypothalamic measurements of POMC, NPY, and AgRP

The hypothalamus was freshly isolated and homogenized in TriZol (Thermo Fisher). Total RNA was further purified using RNeasy column, all following the manufacturer’s instructions. RNA concentration was determined with NanoDrop (Thermo Fisher). One microgram of RNA was used for reverse transcription using a standard method. The resultant cDNA was used to quantify levels of genes of interest (FAM dye) and house-keeping beta-actin (VIC dye) in a single well (input difference was eliminated) using the TaqMan method on a QuantStudio3 machine. TaqMan primers/probes were purchased from Integrated DNA Technology (IDT): Pomc Mm.PT.58.5917321; Npy Mm.PT.58.29444574; Agrp Mm.PT.58.45969609.

### Branched-chain amino acid (BCAA) measurements

A spectrophotometric assay that measures total plasma BCAAs (combined leucine, isoleucine, and valine) was performed as previously described [18]. Briefly, the reaction buffer was first prepared by adding 0.1M glycine, 0.1M KCl, 0.1 KOH (all at pH 10), 0.2M EDTA (pH 8), and double-distilled H2O. The final pH of the solution was kept between 10.5 and 10.7. NAD was dissolved in sodium carbonate buffer (pH 10.7). For 5ul of the sample or standard in each well, 5ul of NAD and 135ul of the reaction buffer were added for the first reading at 340nm. Then, 5ul of leucine dehydrogenase (200U/ml) was added to each well, and the plate was kept undisturbed in the dark for 30 min. The second reading was taken at 340nm, and the absorbance values from the first reading (i.e. baseline) were subtracted from the second reading (i.e. reaction). All samples were run in duplicate within the same assay plate. BCAA concentrations were calculated based on the standard curve equation.

### Body composition measurement

A LF50 mq7.5 nuclear magnetic resonance (NMR) analyzer (Bruker) at the University of Iowa Phenotyping Core Facility was used to measure the fat and lean mass composition of the mice before and after the interventions.

### Histology

Stomach, liver, and small intestine tissue (ileum) were harvested and embedded in an optimal cutting temperature (OCT) compound in a cryomold. Sections 10 µm thick were cut with a cryostat. Sections were stained with hematoxylin & eosin (H&E). For gastric histology, tissue blocks were oriented to obtain sections through the corpus along the greater–lesser curvature axis, including the limiting ridge, fundus (proximal thin-walled compartment rich in ghrelin-producing cells), and body. For small intestinal histology, average villus height/ crypt depth ratio per mouse, was used to assess ileal mucosal thickening. For liver, sections were taken from a standardized central lobe region. For quantitative analyses, gastric mucosal thickness (from muscularis mucosae to luminal surface) and/or fold height was measured in 3–5 non-overlapping fields per section at 20× magnification and averaged to yield a single value per animal.

### Blinding

Surgeons were necessarily aware of group allocation during the procedures. After surgery, animals were coded to keep post-operative investigators blinded. Data sets used for statistical analysis were labeled only with group codes, and group identities were revealed to the analyst only after the primary analyses and figures were finalized.

### Statistical analysis

All reported data are presented as mean ± standard error of means (SEM) with statistical analysis by Student’s t-test, one-way or two-way ANOVA, ANCOVA (with lean mass as covariate for energy expenditure analyses) followed by correction analysis if necessary. Figures and analyses were generated using "GraphPad Prism version 10.6.1 (GraphPad Software, www.graphpad.com)and a p-value of < 0.05 was considered statistically significant. For metabolic phenotyping (n=4–5 per group): Power calculations assumed a moderate effect size (Cohen’s f=0.5 for ANOVA across Sham, SG, GP), standard deviation (SD) of 15–20% from pilot EE/VO₂ data, alpha=0.05, and power=0.80, yielding n=4–5 animals/group to detect group differences in energy expenditure or RER post-intervention.. For molecular/hormone assays (n=8–9 per group): Based on hypothalamic mRNA (e.g., POMC, NPY) or serum leptin data, assuming effect size f=0.4 (from prior SG studies showing 30–50% gene expression changes), SD=25%, yielding n=8/group for >80% power in one-way ANOVA.

## Results

### Comparable Weight Loss but Divergent Body Composition Effects

Both sleeve gastrectomy (SG) and clip-based sleeve gastroplasty (GP) **(Figure 1A)** induced marked weight reduction over six weeks. Following an initial postoperative dip, body weight gradually stabilized. By week six, SG and GP resulted in comparable total weight loss of 12% and 11%, respectively **(Figure 1B, C)**. Body composition analysis using nuclear magnetic resonance (NMR) suggested somewhat different physiological adaptations between the two procedures: SG mice showed a more pronounced reduction in fat mass with a modest increase in lean mass, whereas GP mice exhibited smaller, directionally similar changes that did not reach statistical significance at the current sample size (Figure 1D, E).

**Figure 1.**
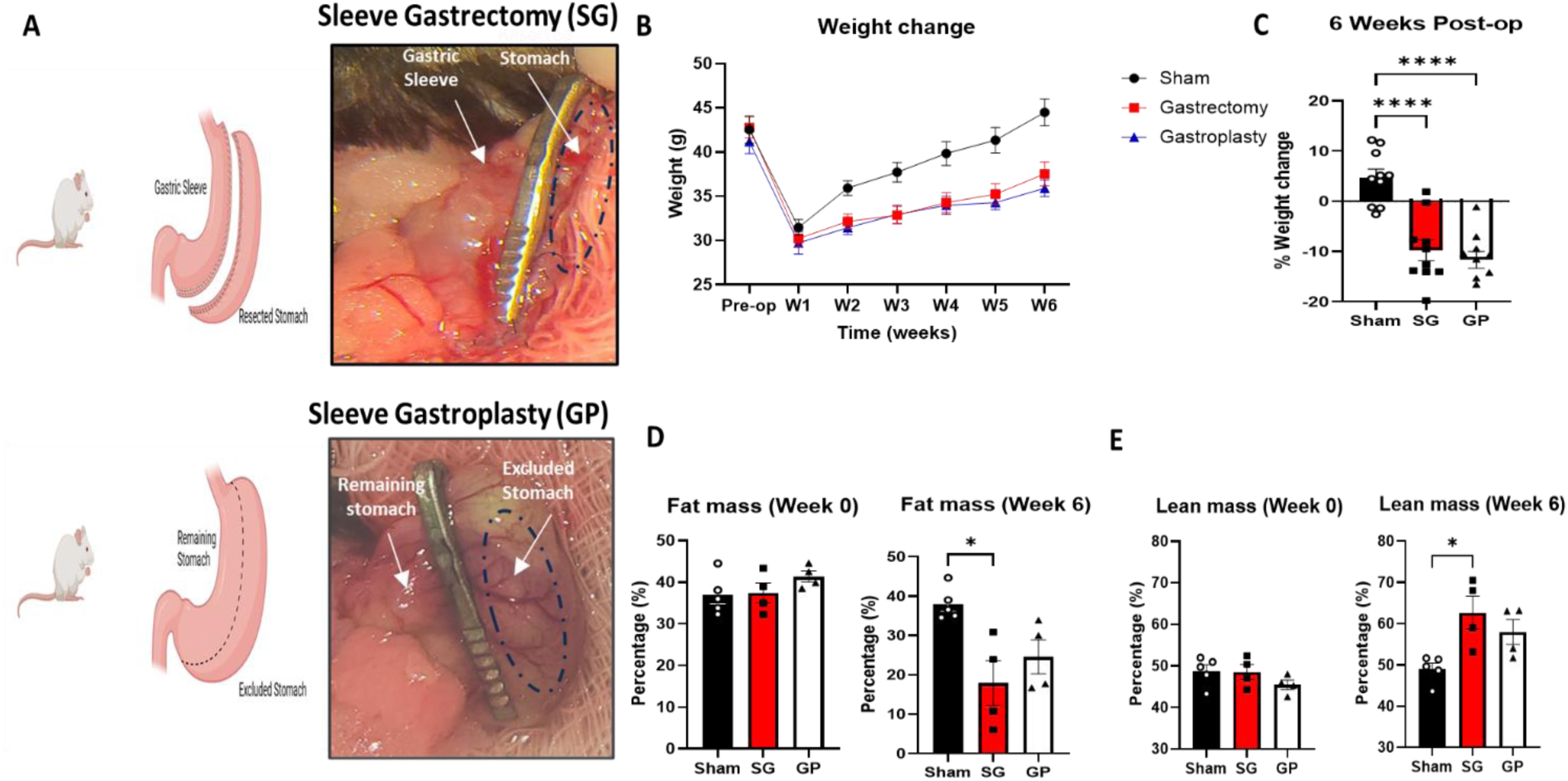
(A) Scheme showing anatomical changes post sleeve gastrectomy (SG) and sleeve gastroplasty (GP). (B) Weight change in (g) over time in (weeks) and (C) % weight change at week 6 post-intervention. Body composition using the NMR LF50 machine: (D) % body fat mass and (E) % lean mass at Week 0 and on day of sacrifice (Week 6). n =8-9 for body weight experiments; n=4-5 for body composition experiments; *p < 0.05, ****p < 0.0001.

Histological examination revealed pronounced gastric mucosal hypertrophy in the SG group **(Figure 2A)**, consistent with local mechanical and regenerative responses to surgical resection. Interestingly, the ileal mucosa remained histologically unaltered across experimental groups **(Figure 2B)**, suggesting that these hypertrophic effects were localized to the gastric compartment rather than representing systemic or pan-enteric changes.

**Figure 2.**
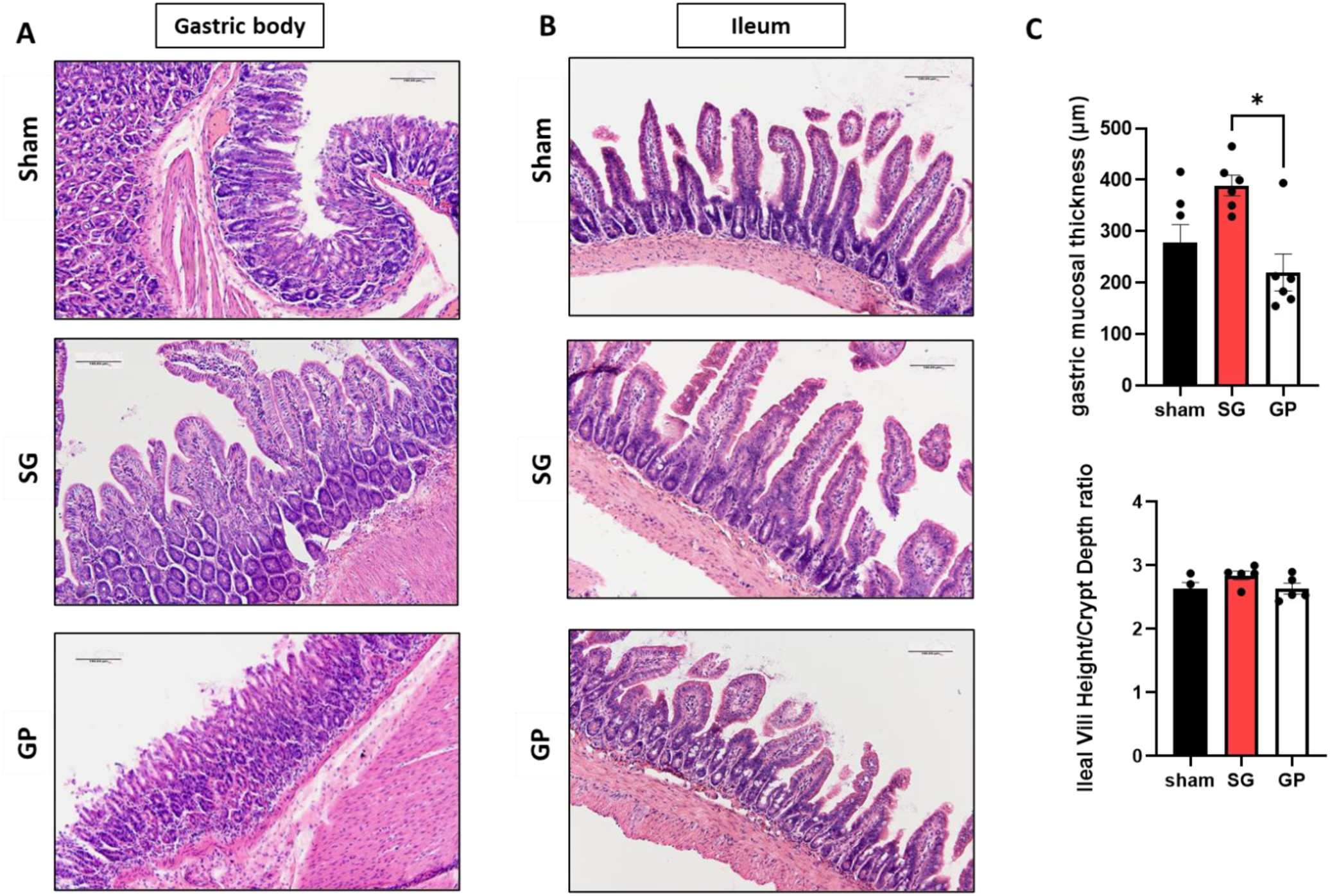
Hematoxylin and eosin (H&E) stain of (A) the gastric body and (B) distal small intestine (ileum) of C57 Blk/6J diet-induced obese male mice that underwent Sham vs Sleeve Gastrectomy (SG) vs Sleeve Gastroplasty (GP) surgery. (C) Average gastric mucosal thickness (expressed in µm) and ileal mucosal thickness (expressed in villi height/crypt depth ratio. All images are at 20x magnification. n=4-5.

### Distinct regulation of energy balance between Sleeve Gastrectomy and Sleeve Gastroplasty

Food intake showed temporally distinct modulation between SG and GP. SG mice displayed a transient yet pronounced reduction in food consumption during week 2 post-surgery, whereas the GP group exhibited a delayed reduction during week 3, with both returning to baseline intake by week 4 **(Figure 3A)**. Notably, SG mice demonstrated altered feeding microstructure characterized by smaller meal sizes and increased meal frequency during week 4 compared to both GP and sham controls **(Figure 3B, C)**, indicating shifts in behavioral regulation rather than simple caloric restrictions. Metabolic cage assessments revealed that, by ANCOVA with lean mass as a covariate, total energy expenditure did not differ significantly among sham, SG, and GP groups. For visualization, TEE normalized to body weight is also shown but statistical inference is based on the lean-mass-adjusted ANCOVA model (Figure 4A). Similarly, average VO₂, VCO₂, and respiratory exchange ratio were comparable across groups (Figure 4B–D). **(Figure 4A–D)**. Hypothalamic profiling uncovered a selective regulatory pattern: POMC expression was significantly reduced in both SG and GP mice relative to sham controls, coinciding with lower circulating fasting leptin levels **(Figure 4E, F)**. However, only SG induced significant reductions in NPY and AgRP expression **(Figure 4G, H)**, pointing to a surgery-specific modulation of central orexigenic pathways. Collectively, these results reveal distinct neuronal signatures driving energy homeostasis after SG and GP.

**Figure 3.**
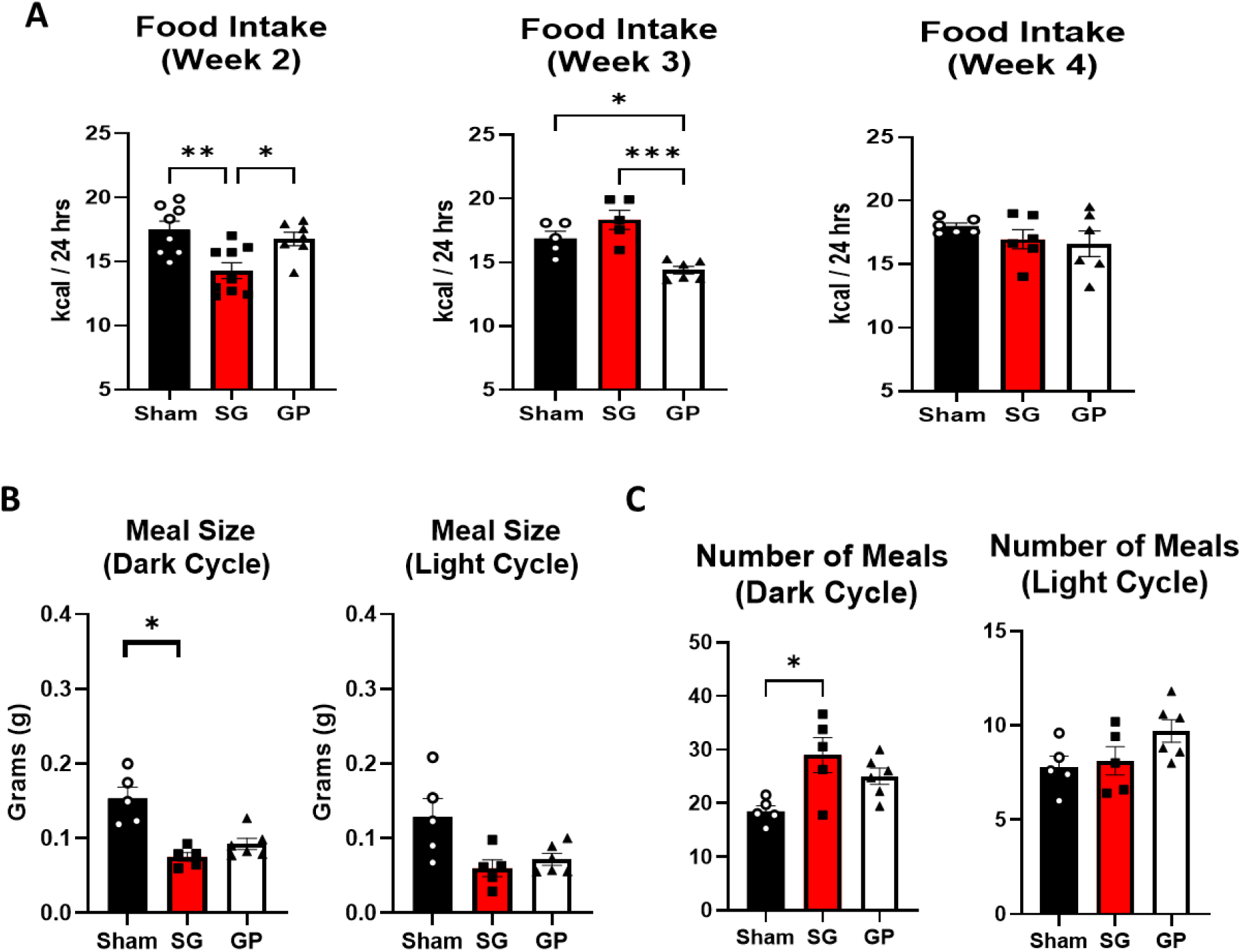
(A) Average daily food intake (kcal/24 hrs) at Week 2, 3, and 4 after Sham, SG, and GP. (B) Average meal size (g) and (C) number of meals during dark and light cycles during week 4 post-intervention. n =5-9 ; *p < 0.05, **p < 0.01.***p < 0.001.

**Figure 4.**
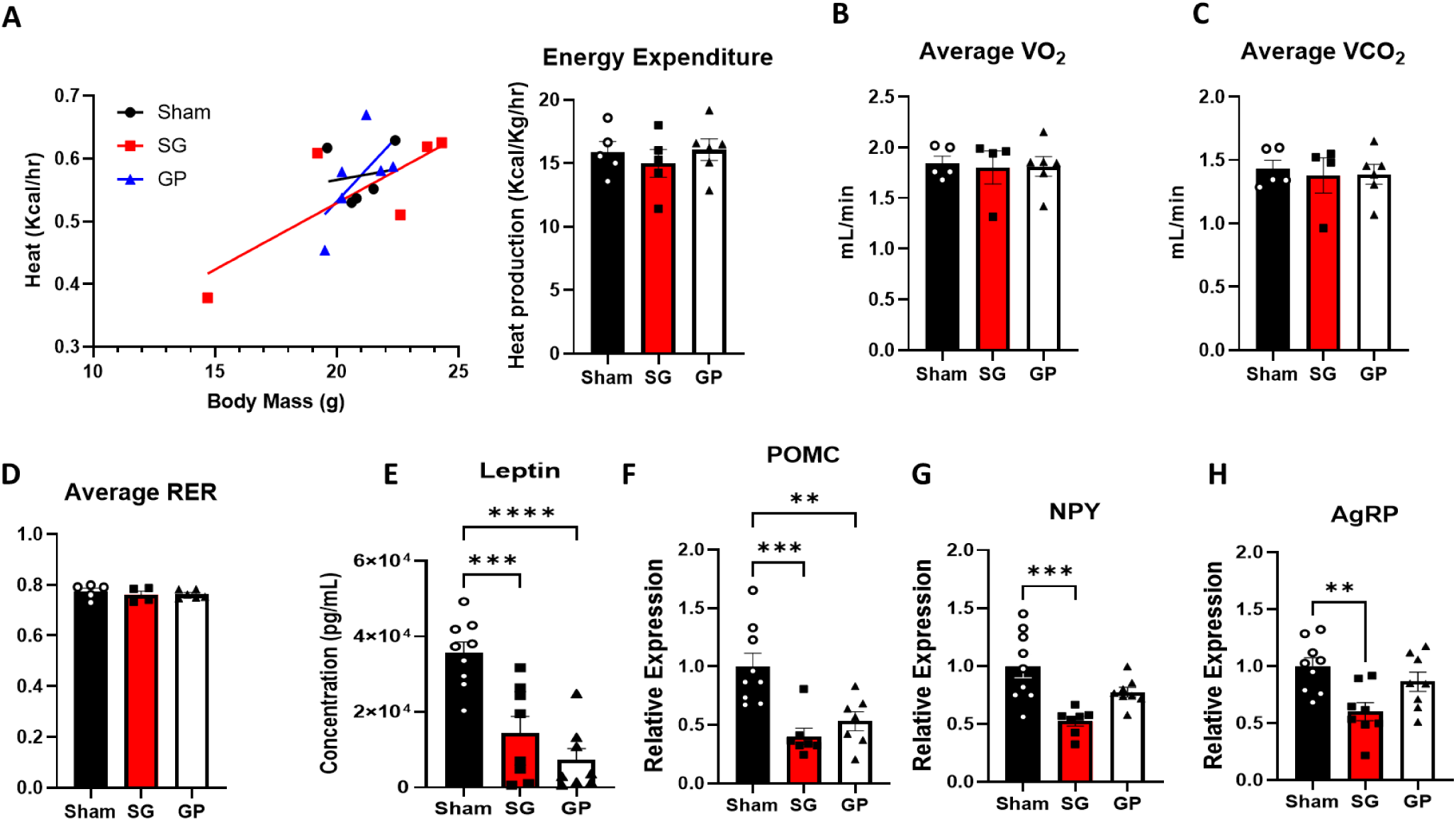
(A) Total energy expenditure (kcal/hr) by indirect calorimetry using Promethion system analyzed by ANCOVA with lean mass as covariate) [*left panel*], and Average energy expenditure (expressed in kcal/Kg/hr) [*right panel*], (B) average oxygen consumption VO2 (mL/min), (C) carbon dioxide production VCO2 (mL/min), and (D) respiratory exchange ratio RER in diet-induced obese C57Bl6J mice at week 4 post Sham versus Sleeve Gastrectomy (SG) versus Sleeve Gastroplasty (GP) intervention. (E) Fasting serum leptin levels and hypothalamic mRNA expression of (F) Proopiomelanocortin (POMC), (G) Neuropeptide Y (NPY), and (H) Agouti-Related Peptide (AgRP) in sham- versus SG- versus GP-operated C57Bl/6J mice at week 6 post-intervention. ANCOVA with lean mass covariate for panel A; one-way ANOVA for remaining panels. n=4–5 (metabolic cage); n=8–9 (mRNA). **p < 0.01, ***p < 0.001, ****p < 0.0001.

### Weight-independent improvement in glucose and lipid metabolism post Sleeve Gastrectomy

Glucose tolerance (GTT) and insulin tolerance tests (ITT) conducted in weeks 4 and 5 postoperatively revealed that both SG and GP improved glucose handling relative to sham controls, though area-under-the-curve (AUC) analysis indicated more pronounced glucose tolerance enhancement in SG **(Figure 5A)**. Given the reported role of branched-chain amino acids (BCAAs) in glucose homeostasis and insulin sensitivity through AgRP-mediated pathways [18] and prior findings suggesting that BCAA reduction post-SG is likely responsible for the post-procedure improvement in glucose metabolism [19], we assessed plasma BCAA levels. SG selectively reduced total circulating BCAAs relative to GP and sham controls **(Figure 5B)**. Although plasma triglycerides (TG) were similar across groups, hepatic TG content, expressed as mg TG per g of liver tissue, was markedly reduced in SG mice compared with sham and GP **(Figure 6A)**. Histological evaluation supported these biochemical findings, showing reduced hepatic steatosis and fewer lipid vacuoles in SG mice **(Figure 6B)**. Collectively, these data indicate that SG confers distinctive, weight-independent metabolic benefits, particularly in glucose regulation and hepatic lipid metabolism

**Figure 5.**
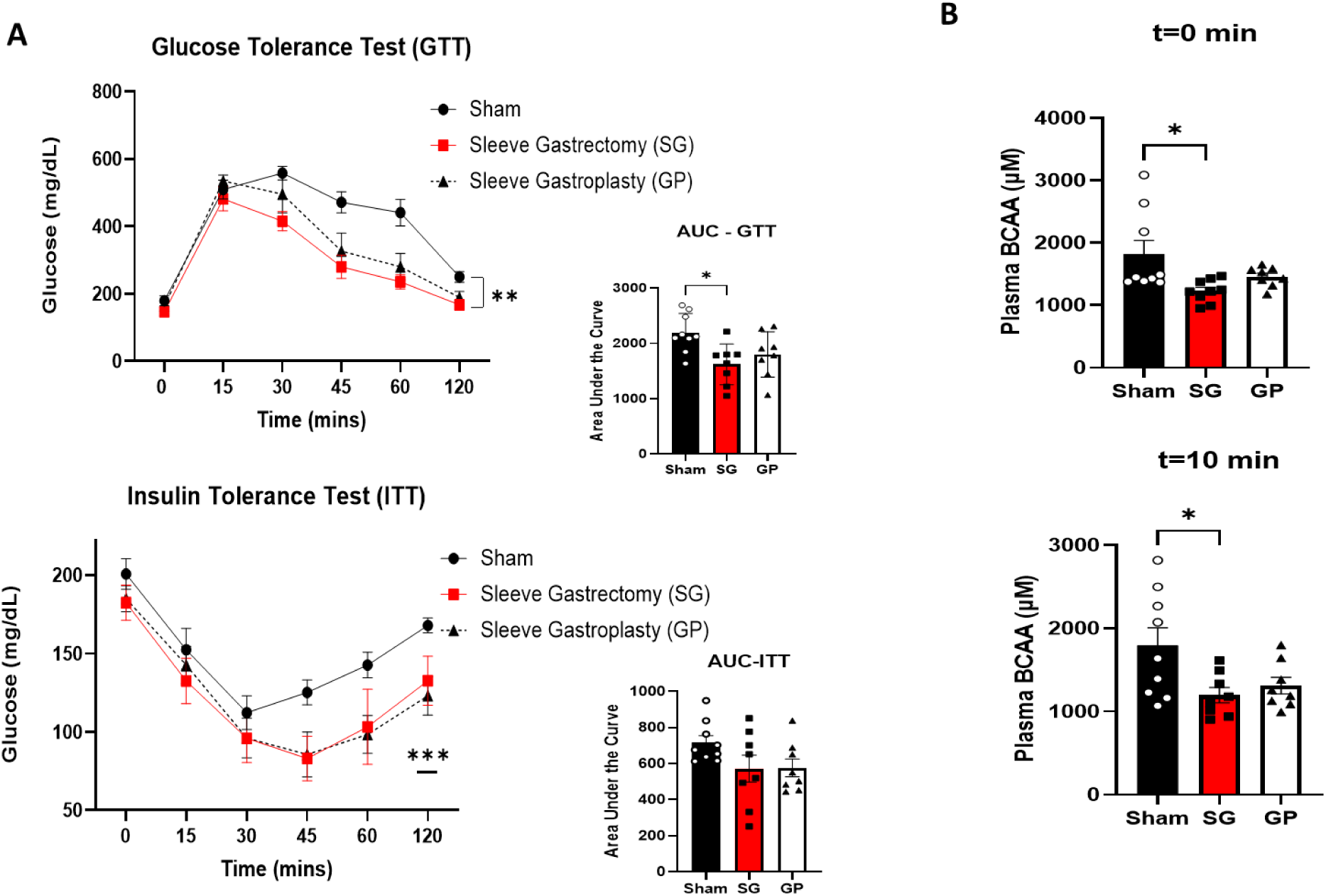
(A) A glucose tolerance test (GTT) and insulin tolerance test (ITT) performed at weeks 4 and 5, respectively, with corresponding area under the curve (AUC) analysis, and (B) plasma branched-chain amino acid (BCAA) levels at time zero (t=0 min) and 10 mins after oral glucose gavage (1g/Kg) at week 6. n=8-9; *p < 0.05, **p < 0.01, ***p < 0.001.

**Figure 6.**
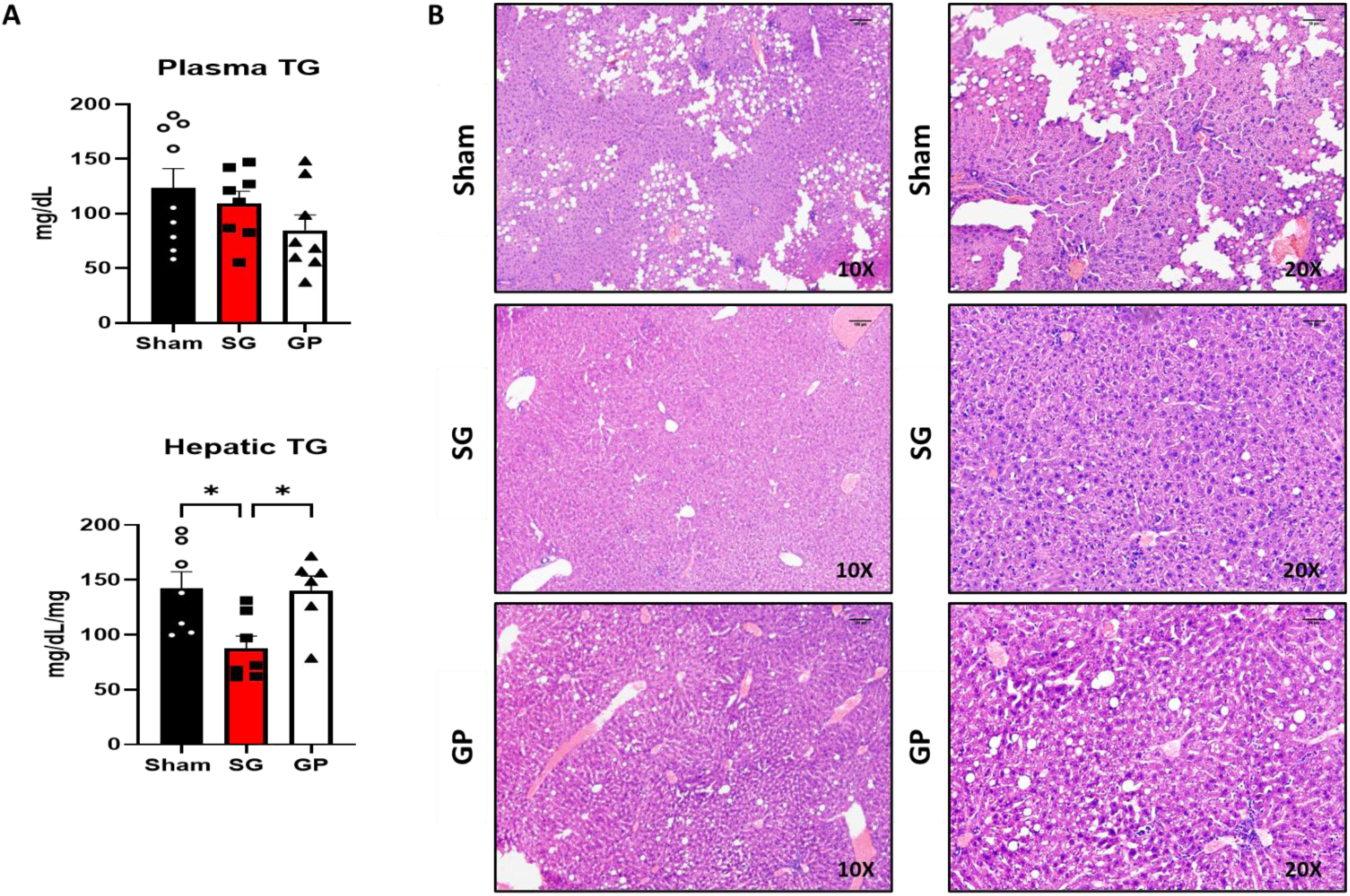
(A) Measured plasma triglyceride (TG) levels (mg/dL) and hepatic TG concentrations normalized by liver (LV) weight (mg/dL/mg) 6 weeks post interventions. (B) Hematoxylin and eosin (H&E) stain of the liver at (A) 10x and 20x magnification; n=6–9.; *p < 0.05.

### Gut Hormonal Profile Post Sleeve Gastrectomy and Gastroplasty

Fasting serum profiling revealed that SG mice exhibited significantly elevated glucagon-like peptide-1 (GLP-1) and peptide YY (PYY) levels, coupled with a reduction in ghrelin concentrations, compared with GP and sham groups **(Figure 7A)**. Fasting C-peptide and insulin were correspondingly decreased in both SG and GP animals relative to their sham counterparts. Following oral glucose administration, two-way ANOVA with factors “group” and “time” demonstrated significant group×time interactions for GLP-1, C-peptide, insulin, glucagon, GIP, and amylin, driven by markedly higher glucose-stimulated levels in SG relative to GP and sham **(Figure 7B)**. These hormonal responses emphasize an exaggerated incretin effect and a broader endocrine adaptation that is unique to SG.

**Figure 7.**
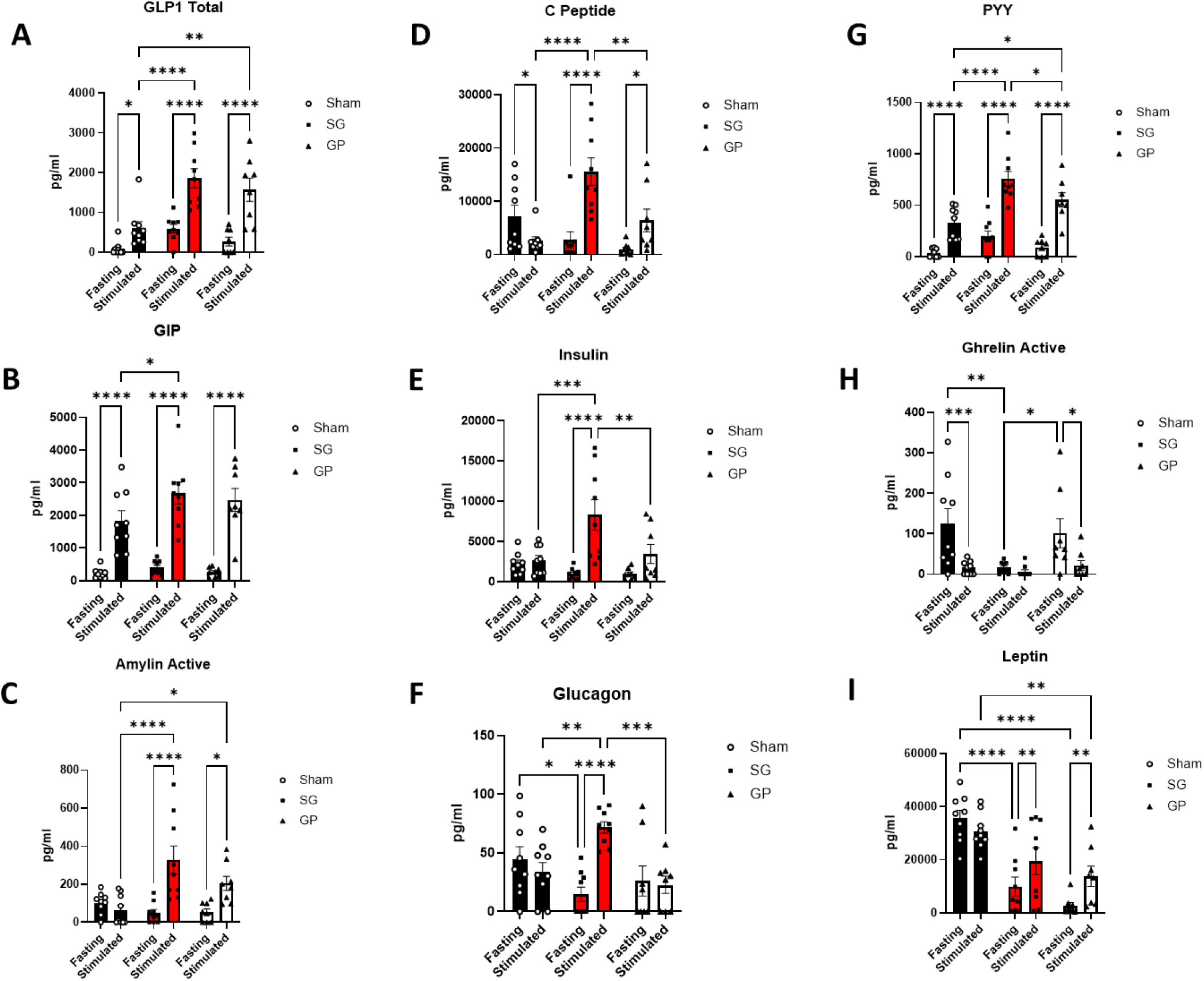
Fasting (time= 0 mins) and stimulated (time=10 mins post-oral glucose gavage) serum levels of (A) Glucagon-like peptide 1 (GLP-1), (B) Glucose-dependent insulinotropic peptide (GIP), (C) Amylin, (D) C-Peptide, (E) Insulin, (F) Glucagon,, (G) Peptide-YY (PPY), (H) Ghrelin, and (I) Leptin in diet-induced obese C57Blk/6J mice 6 weeks after Sleeve Gastrectomy (SG) vs Sleeve Gastroplasty (GP) vs Sham controls. Two-way ANOVA was used to compare means among groups. n =7-9 ; *p < 0.05, **p < 0.01, ***p < 0.001.****p < 0.0001.

## Discussion

Weight loss strategies range from lifestyle interventions, which typically achieve modest reductions of 5–10% in body weight, to pharmacotherapies achieving 8–21%, and bariatric surgeries, which yield sustained 25–30% reduction [20]. In this study, using controlled rodent models of sleeve gastrectomy (SG) and sleeve gastroplasty (GP), both procedures produced comparable weight loss over six weeks but elicited divergent physiological and molecular responses governing energy homeostasis. Clinical evidence parallels our findings. In human cohorts, laparoscopic SG achieves a mean weight reduction of 5.5% within the first month post-operation, compared with 4% in endoscopic sleeve gastroplasty (ESG) [21]. Longer-term follow-ups reveal convergence of outcomes between ESG and SG in terms of weight loss and metabolic disease remission [21,22].

In this study, using controlled rodent models of sleeve gastrectomy (SG) and a surgically created clip-based sleeve gastroplasty (GP), both procedures produced comparable weight loss over six weeks but elicited divergent physiological and molecular responses governing energy homeostasis. Although our rodent GP model does not entirely replicate the human ESG technique since it does not completely exclude the gastric fundus, it remains a valid experimental analogue for comparative investigation. This distinction is essential, as our data confirm similar gross weight changes between SG and GP, yet distinct neurohormonal adaptations. The temporal pattern of food intake reduction, occurring earlier in SG than in GP, suggests differing mechanisms of satiety signaling and feedback. Such transient anorexic responses are consistent with prior rodent studies demonstrating short-lived decreases in intake after bariatric surgery [13]. The persistent alterations in SG food microstructure (smaller, more frequent meals) even after caloric normalization suggest a neuromodulatory reprogramming of appetite circuitry rather than peripheral nutrient sensing. Central to these effects are hormonal and neuronal regulators, particularly NPY, AgRP, and POMC, which influence appetite and energy expenditure. NPY and AgRP are anabolic peptides that stimulate food intake and reduce energy expenditure [22]. This hypothesis is reinforced by our molecular data showing downregulation of hypothalamic orexigenic mediators NPY and AgRP exclusively in SG mice, while both SG and GP exhibited reduced POMC expression, which is likely an adaptation to the newer energy status. The selective modulation of AgRP and NPY aligns with prior studies indicating that specific gastric resection regions, such as the fundus, trigger central appetite reprogramming of hypothalamic circuits likely through vagal or spinal sensory neurons [23].

Beyond weight loss, SG notably improved glucose handling and hepatic steatosis in a manner uncoupled from body weight. These effects reflect findings in both rodent and human studies demonstrating weight-independent metabolic benefits following SG and RYGB [21-23]. Mechanistically, such improvements have been linked to enhanced incretin activity and altered amino acid metabolism. Our observation that SG, but not GP, markedly reduced circulating BCAAs (compared to sham controls) supports data implicating these metabolites in insulin resistance and glucose dysregulation via AgRP neurons [18], consistent with reports of reduced BCAAs post-SG contributing to metabolic improvement [19]. These effects coincide with the observed downregulation of hypothalamic AgRP, suggesting a gut-brain-metabolic feedback axis unique to gastrectomy. The hormonal data further delineate this distinction. SG elicited a pronounced elevation in fasting and glucose-stimulated GLP-1, PYY, GIP, and amylin, surpassing levels seen after GP. GLP-1 and PYY not only enhance insulin secretion and sensitivity but also modulate satiety and gastric motility, thereby amplifying metabolic control [24]. Ghrelin suppression after SG aligns with resection of ghrelin-secreting gastric tissue and may further contribute to appetite attenuation. However, previous studies suggest that effects on food intake, body weight, and glucose tolerance post-gastrectomy in rats are independent of ghrelin [25]. The observed rise in fasting GLP-1 and the exaggerated postprandial incretin response suggest both structural and endocrine remodeling of gut–gut-pancreatic signaling following SG. Human ESG studies consistently show substantial weight loss and improved insulin sensitivity, but gut hormone changes are modest and less consistent than with laparoscopic SG. Fasting ghrelin, GLP-1, and PYY often show no significant early change, with some data suggesting later increases in fasting GLP-1 and PYY at longer follow-up [26, 27]. Similar to our findings in rodent, laparoscopic SG in human subjects reliably lowers ghrelin and significantly increases postprandial GLP-1 and PYY, indicating a stronger endocrine (incretin/orexigenic) component [26]. The selective downregulation of hypothalamic NPY/AgRP expression, together with reduced circulating BCAAs and exaggerated incretin responses, points to a resection-dependent gut–brain–metabolic axis that is not fully engaged by non-resection, exclusion-based gastroplasty. In clinical terms, this suggests that procedures that preserve the gastric fundus and greater-curvature tissue may achieve meaningful but potentially less pronounced neuroendocrine remodeling than SG, consistent with reports of more modest gut-hormone changes after ESG. At the same time, our data should not be over-interpreted as a direct surrogate for ESG, since the GP model differs in access route, device mechanics, and exact gastric geometry.

Cumulatively, these findings reinforce that bariatric procedures exert complex neuroendocrine and metabolic effects extending beyond simple mechanical restriction or malabsorption. SG, in particular, accelerates nutrient transit and alters gut–brain communication, leading to hormone-mediated metabolic benefits [28]. Direct and indirect comparisons between SG and ESG consistently show SG achieving superior weight loss (30.5% total weight loss [%TWL] and 69.3% excess weight loss [%EWL] at 12 months vs. ESG’s 17.1% TWL and 63% EWL), though ESG offers a markedly safer profile with lower adverse events (2.9% overall vs. 11.8% for SG; reduced bleeding [1.1% vs. 2.6%] and GERD [0.4% vs. 5.8%]).[29] Four-year follow-up data on ESG showed that % TWL is ∼ 18% and %EWL is ∼49% [30]. No head-to-head randomized trials are yet available; however, propensity-matched cohorts and MBSAQIP registry data consistently support ESG as a less invasive option with faster recovery, reversibility, and particular applicability in patients with milder obesity (BMI 30–40). These distinctions argue for a tailored approach in surgical and endoscopic obesity management, wherein the choice of intervention aligns with individual metabolic needs.

While ESG offers important advantages in terms of reduced invasiveness, reversibility, and a lower adverse-event profile compared with laparoscopic SG, several cohorts indicate that its long-term weight-loss durability is more modest, with partial weight regain over time and lower total weight loss at multi-year follow-up. These observations suggest that purely restrictive, non-resection approaches may not fully reproduce the durable metabolic effects seen after SG and highlight the importance of dissecting which components of SG biology—such as gastric resection, fundus removal, or more profound gut-hormone changes—are necessary for sustained benefit. Our clip-based GP model captures the non-resection restriction component but, by design, is not an ESG procedure and may underestimate or overestimate certain clinical ESG effects.

### Study Limitation

This work has several limitations. First, the study was conducted in diet-induced obese male C57BL/6J mice with relatively small group sizes, which limits statistical power and means that some apparent differences between SG and GP, particularly for body composition and certain hormonal endpoints, should be interpreted cautiously as hypothesis-generating rather than definitive. Second, our clip-based GP is a surgically created laparotomy-based model that narrows the gastric body while preserving the fundus and greater-curvature segment; it does not replicate endoscopic sleeve gastroplasty in terms of transoral access, endoluminal suturing, or exact gastric geometry. Third, follow-up was limited to six weeks, so we cannot address the long-term durability of weight loss or metabolic changes, which are clinically critical for both SG and ESG. Finally, we focused on selected hormones, hypothalamic transcripts, and BCAAs; broader omics-level profiling and direct functional studies of neural pathways would be needed to fully define the mechanisms underlying the observed resection-specific effects.

In conclusion, our study highlights that both SG and clip-based GP achieve comparable short-term weight loss but diverge substantially in neurohormonal regulation, glucose metabolism, and hepatic lipid handling. SG exerts weight-independent metabolic advantages mediated by altered hypothalamic signaling, reduced circulating BCAAs, and enhanced incretin responses. Within the constraints of this rodent model and its differences from clinical ESG, these findings support the concept that gastric resection adds metabolic value beyond non-resection restriction alone and may help inform the design and optimization of future minimally invasive bariatric and metabolic interventions.

## Author Contribution

Conceptualization: M.J. and M.M.; Data curation: M.J., D.T., S.S, A.F., Y.C., Z.F.H.; Formal Analysis, M.J., D.T., S.S, A.F., Y.C., Z.F.H ; Investigation, M.J., D.T., S.S, A.F., Y.C,Z.F.H, A.C.S., and M.M. Manuscript Writing & Critical Revision: M.J., and M.M. Funding: A.C.S., M.M.

## Funding Sources

M.M. is supported by funds from the US Department of Veterans Affairs, VA Merit Review Program (BX004774-05), the Department of Internal Medicine at the University of Iowa Carver College of Medicine, and the University of Iowa Fraternal Order of Eagles Diabetes Research.

## Declaration of Interests

The authors declare no competing interests.

## Data availability

The datasets used and/or analyzed during the current study are available from the corresponding author on reasonable request.

## Notes

### Competing Interest Statement

The authors have declared no competing interest.

